# SIN3 acts in distinct complexes to regulate the germline transcriptional program in *C. elegans*

**DOI:** 10.1101/2023.03.07.531480

**Authors:** M. Caron, V. Robert, L. Gely, A. Adrait, V. Pakulska, Y. Couté, M. Chevalier, C.G. Riedel, C. Bedet, F. Palladino

## Abstract

The SIN3 transcriptional coregulator influences gene expression through multiple interactions that include histone deacetylases (HDACs). Haploinsufficiency and mutations in SIN3 are the underlying cause of Witteveen-Kolk syndrome and related intellectual disability (ID)/autism syndromes, emphasizing its key role in development. However, little is known about the diversity of its interactions and functions in developmental processes. Here we show that loss of SIN-3, the single SIN3 homologue in *Caenorhabditis elegans*, results in maternal effect sterility associated with deregulation of the germline transcriptome, including desilencing of X-linked genes. We identify at least two distinct SIN3 complexes containing specific HDACs, and show that they differentially contribute to fertility. Single cell smFISH reveals that in *sin-3* mutants, the X chromosome becomes re-expressed prematurely and in a stochastic manner in individual germ cells. Furthermore, we identify histone residues whose acetylation increases in the absence of SIN3. Together, this work provides a powerful framework for the *in vivo* study of SIN3 and associated proteins.

## Introduction

The highly conserved SIN3 transcriptional coregulator acts as a scaffold to assemble distinct complexes containing histone deacetylases (HDACs), chromatin adaptors and transcription factors that modify chromatin to influence gene expression (Laherty et al., 1997; Banks et al., 2018; Banks et al., 2020). SIN3/HDAC complexes regulate essential cellular processes, including differentiation, cell cycle regulation and metabolism through both gene activation and repression (Saha et al., 2016; van Oevelen et al., 2010). Importantly, heterozygous loss-of-function variants and point mutations in the two mammalian homologs *SIN3A* and *SIN3B* were recently identified as the underlying cause of Witteveen-Kolk syndrome and related intellectual disability (ID)/autism disorders (Balasubramanian et al., 2021; Latypova et al., 2021; Latypova et al., 2021; Witteveen et al., 2016) , emphasizing the functional importance of these proteins.

In different species SIN3 proteins have been shown to reside in multiple complexes. In yeast these are named Rpd3 large (Rpd3L: Sin3, Rpd3/HDA1, Ume1, Sap30, Sds3/SUDS3, and other proteins) and Rpd3 small (Rpd3S: Sin3, Rpd3/HDA1, Ume1, Rco1 and Eaf3) (Carrozza et al., 2005a; Carrozza et al., 2005b; Kadosh and Struhl, 1997; Kasten et al., 1997). In mammals two highly related proteins, SIN3A and SIN3B, share overall domain structure and association with HDAC1 and HDAC2. Biochemical data from numerous studies suggests that SIN3A and SIN3B function in distinct complexes, and that paralog identity influences complex composition, with SIN3B copurifying in a complex equivalent to yeast Sin3S/Rpd3S (Jelinic et al., 2011; Vermeulen et al., 2010; Adams et al., 2020), and SIN3A residing in multiple larger complexes related to SIN3L/Rpd3L (Banks et al. 2020, 2018; Streubel et al. 2017; Saunders et al. 2017; Fleischer et al. 2003; Alland et al. 2002; Zhang et al. 1997; Adams et al. 2020). Both homologues interact directly with transcription factors through one of three paired-amphipathic helix (PAH) domains, and are recruited to chromatin through accessory proteins including ARID4A/B, PHF12/PF1 and MRG15. Mouse knockout studies have shown that SIN3A and SIN3B are non-redundant (Dannenberg et al., 2005; David et al., 2008), consistent with at least partially distinct functions. However, the presence of two homologues, the temporal and cell type-specific nature of SIN3 complex activities, and the transient association of SIN3 with numerous transcription factors and accessory proteins has hampered the study of individual complexes in a developmental context.

*Caenorhabditis elegans* contains a single SIN3 homologue, SIN-3, facilitating its study (Choy et al., 2007; Beurton et al., 2019). Genetic analysis of *sin-3* mutant animals carrying the molecularly uncharacterized *tm1276* allele revealed roles in male development (Choy et al., 2007), motility, longevity and fertility (Pandey et al., 2018; Sharma et al., 2018), but the specific function of SIN-3 in these different processes remains mostly unknown, as does the identity of its interaction partners. Here we combined genetic analysis with affinity purification coupled to mass spectrometry (MS)-based proteomics, transcriptomics and single cell analysis to more clearly dissect the function of SIN-3. Using a *sin-3* null mutant constructed by CRISPR-Cas9, we uncovered an essential requirement for SIN-3 in germ stem cell proliferation and fertility. Immunoprecipitation followed by MS-based proteomic (IP-MS) analysis confirmed the identity of the previously described SIN3 small (SIN3S) complex containing MRG-1/MRG15, HDA-1/HDAC and ATHP-1/PHF12 (Beurton et al., 2019). In addition, we identified counterparts of known mammalian SIN3L complex subunits including SUDS-3/SDS3 and ARID-1/ARID4, thereby defining at least two distinct SIN3 complexes. We also provide evidence that specific HDACs reside in different SIN3 complexes, and that these differentially contribute to germline health. Genome-wide transcriptomics analysis combined with single molecule fluorescence *in situ* hybridization (smFISH) reveals that loss of SIN-3 results in desilencing of X-linked genes in a stochastic manner, and histone post-translational modification analysis by MS-based proteomics identifies histone H3K18AcK23Ac as a target of SIN-3 dependent deacetylation. Together, our results reveal an essential role for SIN-3 in preserving the germline transcriptional program and fertility, and provide insight on how, within a single tissue, distinct SIN3 complexes and interaction partners contribute to specific regulatory functions.

## Results

### Loss of *sin-3* results in maternal effect sterility

An mCherry::SIN-3 translational fusion protein constructed by CRISPR-Cas9 is ubiquitously expressed in nuclei of both the germline and soma, and in embryos starting at the 4-cell stage ( Figure 1A and S1A). Previous studies using the *sin-3*(*tm1276*) allele revealed defects in the male tail, decreased lifespan, and reduced fertility (Beurton et al., 2019; Choy et al., 2007; Choy et al., 2007; Sharma et al., 2018). Because *tm1276* is a small internal deletion that only removes exon 2 of *sin-3* (Figure 1B), we used CRISPR-Cas9 genome editing to construct a full knock-out allele, *syb2172*, in which the ATG codon and the entire *sin-3* coding region are removed (Figure 1B). *syb2172* homozygous animals were maintained as heterozygotes using a balancer chromosome (see Material and Methods). We observed that F1 homozygous *syb2172* progeny derived from heterozygous mothers, which inherit maternal *sin-3*(+) product but do not synthesize zygotic product (abbreviated M+Z-), produce significant fewer F2 offspring than wild type (mean 120 versus 350, Figure 1C). Second generation F2 animals without maternal contribution (M-Z-) develop into fully sterile adults, with a few animals producing 10 or fewer progeny. *sin-3(tm1276)* animals can instead be maintained as homozygotes, although as previously reported they produce fewer progeny (Beurton et al., 2019; Pandey et al., 2018) (Figure 1C). Together these results show that complete loss of *sin-3* causes sterility in absence of maternal contribution, and indicate that the original *tm1276* mutation is most likely a strong loss-of-function allele that retains partial function.

**Figure 1.**
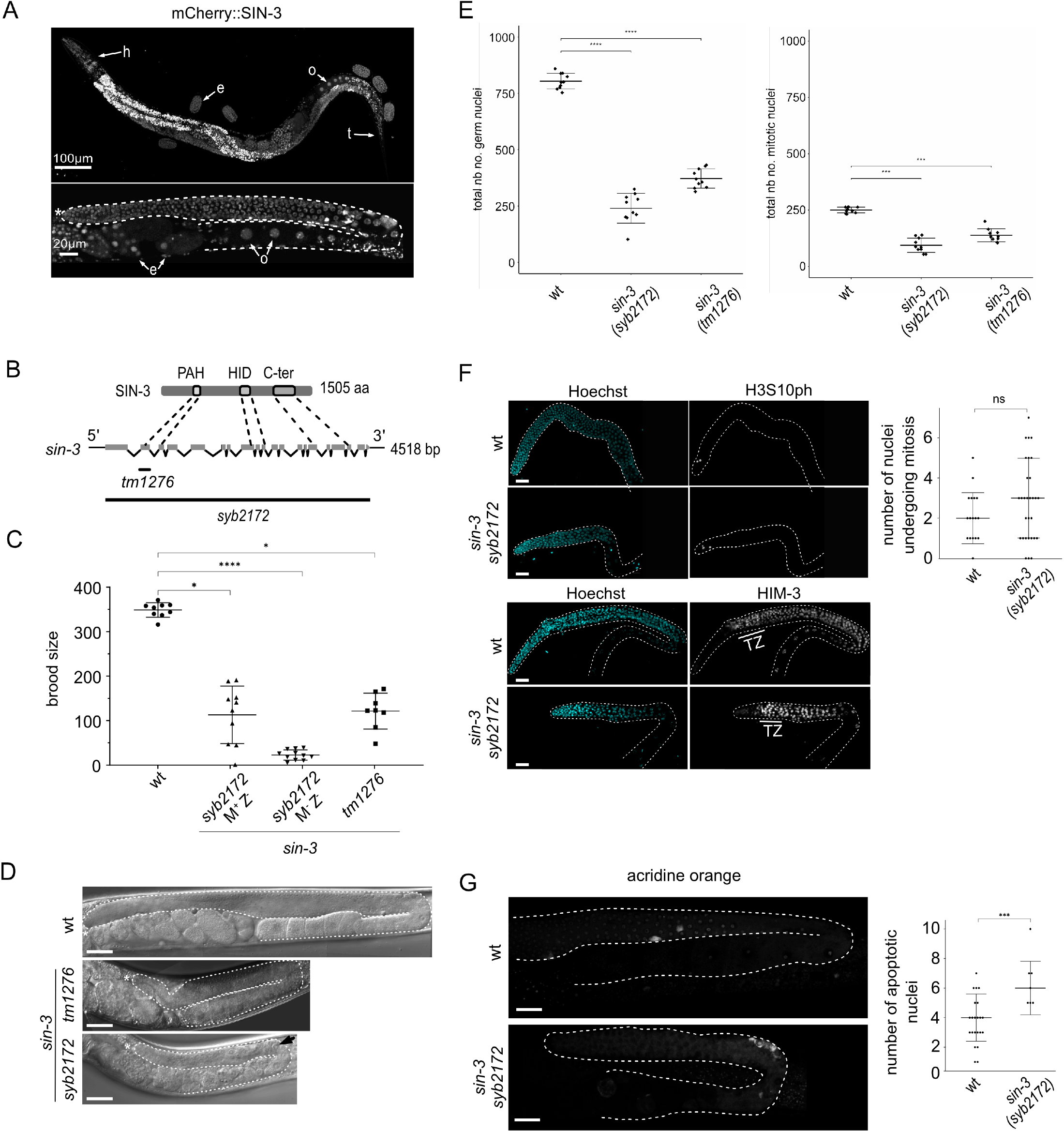
Sterility and reduced progenitor cell number in *sin-3* deletion mutants. (A) Representative image of whole animal expressing endogenously tagged mcherry::SIN-3. Top panel: expression of SIN-3 throughout the body including nuclei in head (h) and tail (t), embryos (e) and oocytes (o). Final image was obtained by stitching each image to its neighbors with the best alignment. Bottom panel: SIN-3 expression throughout the adult germline (one arm of the gonad is shown, outlined with dotted line). (*) marks the distal end of the germline, oocytes (o). Scale bar is 100 µm for whole animal, 20 µm for gonad arm. Images were taken from different animals. (B) Cartoon representation of the SIN-3 protein with conserved domains and the corresponding exon sequence on the *sin-3* gene. The extent of the *tm1276* and *syb2172* deletions are indicated below the gene sequence. (C) Brood size count of wildtype and *sin-3* mutants. M+Z-: homozygous mutant with maternal contribution; M-Z-: homozygous mutant without maternal contribution. Multiple comparison was done using Dunn’s multiple comparisons test following a significant Kruskal Wallis test; * p < 0.05; **** p < 0.0001. (D) Representative differential interference contrast (DIC) microscopy images of germlines from wildtype and *sin-3* mutant animals (M-Z-). Germlines are outlined with dotted line, (*) marks the distal end of the germline. Scale bar is 10µm. (E) Total number of germline nuclei, and number of mitotic nuclei per gonad arm in wildtype and *sin-3* mutant worms. 10 gonad arms were counted per genotype. Multiple comparison was done using a generalized linear model with a quasi-Poisson law. **** p < 0.0001. (F) Representative confocal microscopy images of phospho-H3 (upper panel) and HIM-3 (lower panel) staining on dissected germlines (outlined with dotted lines) from wildtype and *sin-3 (syb2172)* animals. For HIM-3 staining, the transition zone (TZ) defined by Hoechst staining is displayed. Statistical comparison was done using Mann-Whiney-Wilcoxon test. N = 17 for wt and n = 30 for *sin-3(syb2172).* (G) Acridine orange (AO) staining of apoptotic cells in a single gonad arm from whole animals. Dot plot displays the number of apoptotic bodies stained with AO per germline. Statistical comparison was done using Student’s t-test. n = 22 for wt and n = 7 for *sin-3(syb2172)*. *** p < 0.001. Scale bars are 10 µm in F and 15 µm in G.

### Reduced proliferation of progenitor cells in *sin-3* mutant germlines

In the *C. elegans* germline, meiotic nuclei are arranged in a spatio-temporal order, with the distal end of the gonad containing mitotically proliferating nuclei. Subsequent stages, clearly recognizable by DAPI-staining, consist of a “transition zone” where homologue pairing occurs and chromosomes become polarized in a crescent shape, followed by the pachytene stage during which synapsed chromosomes appear as discrete, parallel tracks. More proximally, nuclei exit pachytene, enter diplotene, and cellularized oocytes containing condensed homologues are formed (Crittenden et al., 2006). Adult germlines of *sin-3* mutant animals are significantly smaller than wild type, with a more severe phenotype observed in *sin-3(syb2172)* compared to *sin-3(tm1276)* (Figure 1D). While the overall organization of *tm1276* germlines was similar to wild type, with recognizable germ cells in the distal region and oocytes proximally, numerous abnormalities were observed in F2 M-Z-*sin-3(syb2172)* germlines. These included large cells resembling oocytes appearing precociously in the bend region, and a highly disorganized proximal region (Figure 1D). In approximately 20% of these animals we also observed a ‘Gogo’ phenotype, in which germ cells transition from oocytes to pachytene germ cells and back to oocytes (germ line-oocyte-germ line-oocyte) (Eberhard et al., 2013; Sendoel et al., 2019) (Figure S1B).

Counting of DAPI stained progenitor cells in the distal germline, recognized by their morphology (Crittenden et al., 2006), revealed that in both *tm1276* and s*yb2172* mutants germ cell proliferation was severely reduced: the total number of mitotic nuclei decreased from approximately 250 per gonad arm in wild type to 138 in *tm1276* and less than 100 in *syb2172* mutants. A similar decrease was observed when counting the total number of germline nuclei (Figure 1E). Staining of *sin-3(syb2172)* mutant germlines with antibodies directed against histone H3 phosphorylated on Ser10 (H3S10ph) to mark dividing cells, and HIM-3 to mark meiotic cells (Zetka et al., 1999), revealed no significant difference compared to wild type either in the number of mitotic figures in the distal region, nor in HIM-3 staining beginning at the transition zone (Figure 1F). These results suggest that *sin-3(syb2172)* germlines retain an overall distal-proximal organization similar to wild type. Because phospho-H3 positive cells are in metaphase, a decrease in the number of proliferating germ cells in *sin-3* mutants without a corresponding reduction in the number of mitotic figures may stem from an extended block or pause in metaphase (Golden et al., 2000). In *sin-3*(*syb2172*) germlines we further observed a small increase in apoptosis (Figure 1G). Altogether, these results suggest that reduced fertility in the absence of *sin-3* can be largely accounted for by a decrease in germ cell proliferation, in combination with increased apoptosis. In addition, the developmental switch regulating the transition from late pachytene to oogenesis may be compromised in some animals, as revealed by the “Gogo” phenotype.

### SIN-3 resides in distinct complexes

SIN-3 does not have DNA binding or enzymatic activities on its own, and its co-regulatory functions are dependent on the interaction with its protein partners (Kadamb et al., 2013; Adams et al., 2018). To gain insight in the repertoire of proteins that interact with SIN-3 we performed IP-MS on mCherry::SIN-3-expressing embryos (Figure S2A). SIN-3 co-precipitated counterparts of yeast and mammalian Rpd3S/SIN3S subunits ATHP-1/PHF12, MRG-1/MRG15 and HDA-1 (Rundlett et al., 1996; Carrozza et al., 2005b; Jelinic et al., 2011), and COMPASS-targeting subunit CFP-1, as expected (Figure 2A and Table S1)(Beurton et al., 2019). In addition, we identified counterparts of conserved mammalian SIN3L complex subunit SUDS-3/SDS3, and the ARID4 homologue ARID-1 (Banks et al., 2020; Adams et al., 2020; Banks et al., 2018). Low abundance peptides corresponding to RBA-1 and LIN-53, the two homologues of SIN3L complex component RBBP4/7, where also detected, while peptides corresponding to the distantly related SAP30 homologue Y67D2.7 5 were not present in our IP-MS analysis (Table S1). Other top hits include HDA-3, a second class I HDAC (Shi and Mello, 1998), the histone chaperone NAP-1, previously identified as a SIN3 interactor in drosophila (Moshkin et al., 2009), and the uncharacterized protein C01G6.5 encoding an orthologue of the mammalian PHD finger/forkhead transcription factor TCF19, a known interactor of the NuRD HDAC complex (Mondal et al., 2020; Sen et al., 2017). Additional IP-MS experiments using as bait SIN3L complex subunit ARID-1 tagged with GFP (ARID-1::GFP) (Figure S2A and B) confirmed the presence of SIN-3 in a distinct larger complex containing SUDS-3, HDA-3, HDA-1, C01G6.5 and NAP-1 in addition to ARID-1 (Figure 2A and Table S1), while peptides corresponding to SIN3S complex component ATHP-1 were absent. A number of nematode specific proteins were also identified in both IP-MS experiments (Figure 2A).

**Figure 2.**
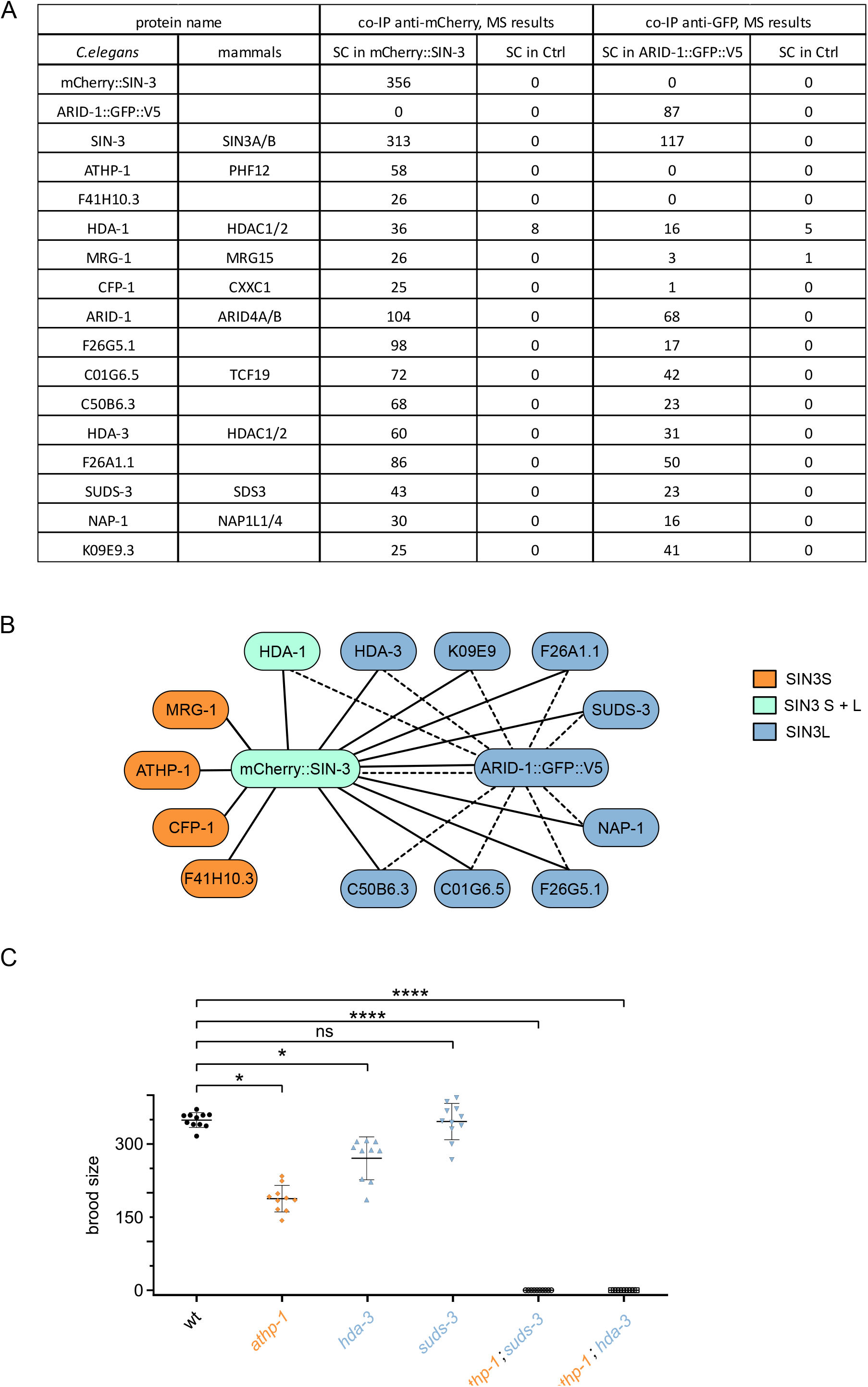
Identification of two SIN-3 related complexes in *C. elegans* embryos by IP-MS, and impact of inactivation of conserved subunits on fertility. (A) List of proteins identified by MS-based proteomics as enriched in mCherry::SIN-3 or ARID-1::GFP compared to the corresponding negative control IP, and identity of their mammalian homologues (full list of identified proteins is provided in Table S1); SC: Spectral Count. (B) Cartoon representation of the protein interactions of SIN-3 and ARID-1, and the complexes inferred from them, based on this data and (Beurton et al., 2019). Subunits of the SIN3S and SIN3L complexes are colored in orange and dark blue, respectively, common subunits are in light blue. (C) Brood size as a read-out of fertility following inactivation of SIN3L and SIN3S complex subunits. Multiple comparison was done using Dunn’s multiple comparisons test following a significant Kruskal Wallis test; ns= not significant ; * p < 0.05 ; **** p < 0.0001.

The above experiments, together with previous data using CFP-1 or MRG-1 as bait in IP-MS experiments (Beurton et al., 2019; Baytek et al., 2022), confirm the presence of SIN-3 in a smaller complex related to yeast and mammalian Rpd3/SIN3S (Carrozza et al., 2005b; Jelinic et al., 2011) containing ATHP-1/PHF12, MRG-1/MRG15, CFP-1 and HDA-1 as the only HDAC. In addition we identify a second complex that we will refer to as SIN3L and that contains SUDS-3, HDA-3, HDA-1, ARID-1/ARID4, NAP-1 and the transcription factor C01G6.5 (Figure 2B). Based on homology with the human protein, we will hereafter designate C01G6.5 as TCF-19. The absence of COMPASS subunits other than CFP-1 from our list of SIN-3 and ARID-1 interactors confirms the presence of CFP-1 independently of COMPASS in a SIN3S related complex (Beurton et al., 2019). Our data also suggest that class I HDACs may differentially contribute to the activity of SIN3 complexes: HDA-1 alone in the SIN3S complex, and HDA-3 together with HDA-1 in a SIN3L complex.

### SIN3L and SIN3S complexes both contribute to germline maintenance

To investigate whether any of the SIN-3 interactors we identified are also required for fertility, we measured brood sizes in the corresponding mutants. SIN3S complex components MRG-1/MRG-15 and HDA-1/HDAC are ubiquitously expressed, well characterized proteins found in additional chromatin complexes and required for fertility and larval development, respectively (Hajduskova et al., 2018; Bleuyard et al., 2017; Iwamori et al., 2016; Iwamori et al., 2016; Smith et al., 2013; Dufourcq et al., 2002; Passannante et al., 2010; Whetstine et al., 2005). Their role in the germline was therefore not further analyzed in this study. We instead focused on one other SIN3S component, ATHP-1/PHF12, and SIN3L complex components HDA-3 and SUDS-3. *athp-1(tm4223)* and *hda-3(ok1991)* are previously described loss of function alleles (Beurton et al., 2019; Kawamura and Maruyama, 2020), while *suds-3(syb2212)* is a full deletion allele constructed by CRISPR-Cas9 for this study. As previously reported, *athp-1* mutants laid significantly fewer progeny than wild type (Beurton et al., 2019)(Figure 2C). A smaller but significant reduction in brood size was also observed in *hda-3* mutants, while *suds-3(syb2212)* mutant animals were fully fertile (Figure 2C). DIC imaging revealed no obvious defects that could account for the reduced brood size of *athp-1*, *hda-3,* or *suds-3* mutants (Figure S3). Because SUDS-3 is a core component of the SIN3L complex and is required for its HDAC activity (Alland et al., 2002b; Lechner et al., 2000), the absence of an obvious germline phenotype in the corresponding mutant suggests that this complex does not play a prominent role in the maintenance of fertility. ATHP-1 is instead an accessory factor in SIN3S complexes whose knock-down may negatively impact SIN3S activity in the germline (Xie et al., 2012; Jelinic et al., 2011; Li et al., 2007). Interestingly simultaneous inactivation of SIN3S and SIN3L complex subunits in both *athp-1;suds-3* and *athp-1;hda-3* double mutant animals resulted in fully penetrant sterility (Figure 2C), suggesting redundant functions in the maintenance of germline function. Alternatively, or in addition, *athp-1*, *hda-3* and *suds-3* may play redundant, *sin-3* independent functions required for fertility (Barnes et al., 2018).

### SIN-3 and HDA-3 have both unique and common regulatory functions in germline gene expression

To explore how the germline transcriptome is affected by loss of SIN-3, we performed transcriptome profiling on dissected gonads from *sin-3(syb2172*) and *sin-3(tm1276)* young adults. We also included *hda-3(ok1991)* mutant germlines in our analysis since HDA-3 is poorly characterized and found in SIN3L but not SIN3S; its transcription profiling could therefore provide useful insight on SIN3L complex function (Figure 3). Using DESeq2 (FDR<0.05) to derive lists of differentially expressed genes, we found a significant number of both up- and down-regulated genes for *sin-3(syb2172*) as compared to wild-type (405 and 683, respectively) (Figure 3A and Table S2), in agreement with SIN-3 acting in both gene repression and activation, as in other systems (Saha et al., 2016; Saunders et al., 2017; Yao et al., 2017; van Oevelen et al., 2008; Gajan et al., 2016). Significantly more genes were found to be misregulated in *sin-3(tm1276)* mutants (2402 up and 3106 down), although there was a large degree of overlap for both up-and dowregulated genes (Figure S4 and Table S2). The difference in the total number of misregulated genes may be attributed to the specific allele used, or to differences in how mutants were propagated: *sin-3(tm1276)* animals were maintained as homozygotes over many generations, which could result in progressive transcriptional deregulation, while *sin-3(syb2172*) animals were maintained as balanced heterozygotes. All subsequent analyses were carried out on the list of genes misregulated in the *sin-3(syb2172*) null allele.

**Figure 3.**
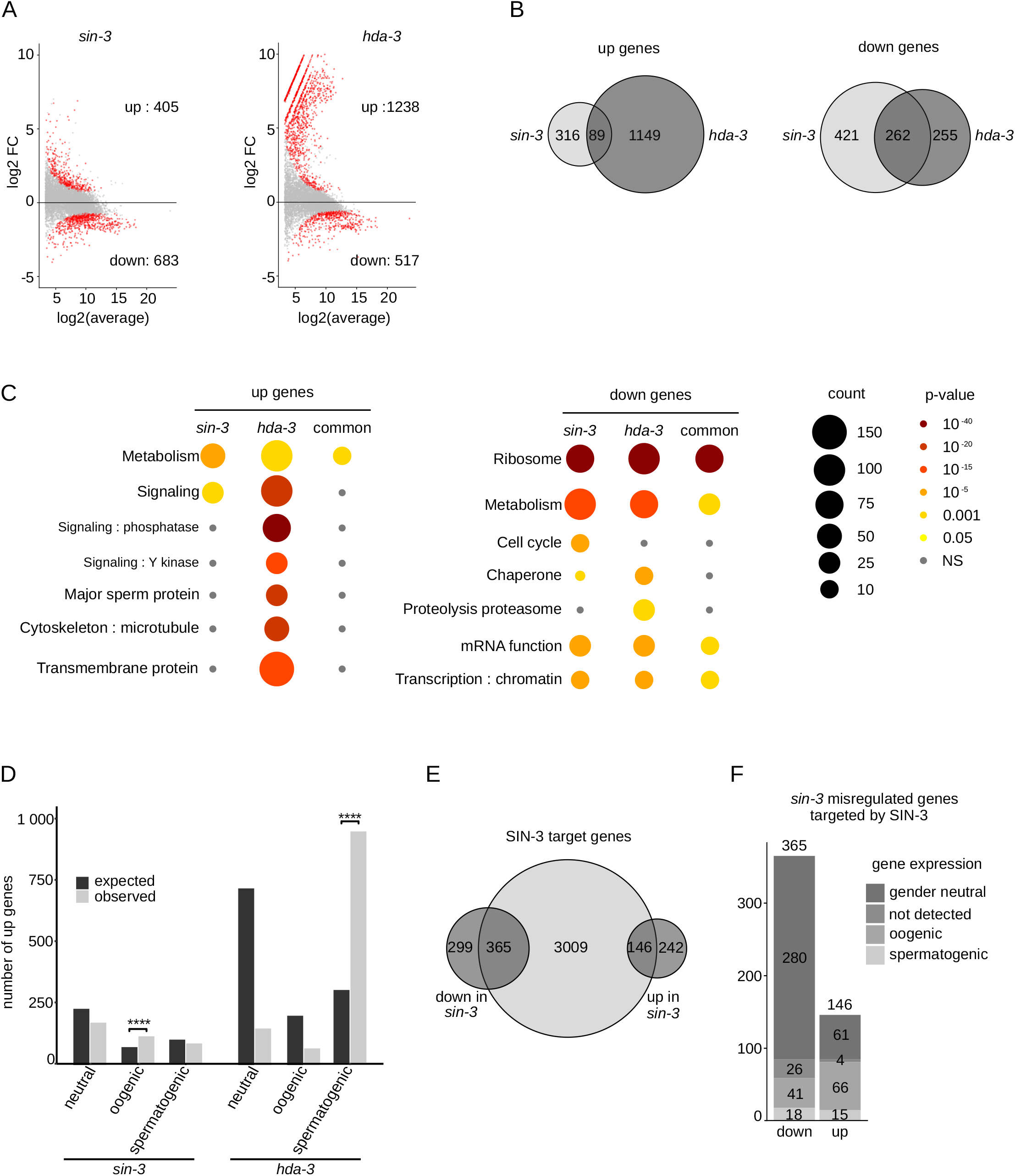
Transcriptomic analysis of *sin-3* and *hda-3* mutant germlines. (A) MA plots showing the log2(fold change) of gene expression in *sin-3(syb2172)* mutants (left panel) and *hda-3(ok1991)* mutants (right panel) compared to wild type as a function of log2(basemean expression level). Genes that are significantly misregulated in each mutant are colored in red and were identified using DESeq2 (with FDR < 0.05). (B) Venn Diagramm showing overlap between upregulated (left) and downregulated (right) genes in *sin-3* and *hda-3* mutants. (C) WormCat visualization of categories enriched in genes upregulated (left) or downregulated (right) in *sin-3* mutants, *hda-3* mutants or common to both (common). The legend for bubble charts is indicated on the right, with size referring to the number of genes in each category and color referring to the p-value. (D) Enrichment analysis. Barplot representing the expected (black) and observed (gray) number of upregulated genes in *sin-3* or *hda-3* mutants present in gene sets derived from transcriptomics of adult oogenic or spermatogenic gonads (Ortiz et al., 2014). Enrichment was calculated with hypergeometric tests performed in R. *** p-value <10^-8^. Gene numbers in each set are the following : total number of protein coding genes 10063 (5950 gender-neutral, 1660 oogenic, 2453 spermatogenic); number of oogenic genes upregulated in *sin-3* (108), in *hda-3* (68); number of spermatogenic genes upreregulated in *sin-3* (79), in *hda-3* (944), number of gender-neutral upregulated genes *sin-3* (173), *hda-3*(149). (E) Venn diagram showing overlap between SIN-3 ChIP peaks on promoters (Beurton et al., 2019) and up- or downregulated genes in *sin-3* mutant germlines. (F) Bar plot representing the repartition of SIN-3 target genes that are up- or downregulated in the different gene sets defined by (Ortiz et al., 2014).

In contast to *sin-3* mutants, in *hda-3* mutants 70% of deregulated genes were up-regulated (1238 out of 1755 total), and of these, only 89 were also upregulated in *sin-3(syb2172)* mutants (Figure 3B). Therefore, as observed for other HDACs (Kadosh and Struhl, 1997; Rundlett et al., 1998; Taunton et al., 1996), *hda-3* acts mainly as a transcriptional repressor, and this function is mostly independent of *sin-3*. Surprisingly instead, over 50% of *hda-3* downregulated genes are also down-regulated in *sin-3(syb2172*) (262 out of 517 total) (Figure 3B). These may represent direct targets of SIN-3/HDAC mediated activation (Wang et al., 2009; Chou et al., 2011; Greer et al., 2015; Kim et al., 2013; Kelly and Cowley, 2013), or indirect targets. RT-qPCR analysis confirmed the differential expression of both up- and down-regulated genes in *sin-3* mutant animals as compared to wild-type (Figure S5). Additional RT-qPCR analysis on *hda-3*, *suds-3* and *athp-1* mutants revealed that 3/5 selected genes downregulated in *sin-3* and *hda-3* were also downregulated in *suds-3* mutants, while their deregulation varied in *athp-1* mutants. For upregulated genes instead, no consistent pattern was observed, although *hda-3* and *suds-3* expression profiles were very similar. This limited analysis is consistent with common deregulation in SIN3L complex mutants.

Enrichment analysis using WormCat (Holdorf et al., 2020) revealed that among downregulated genes, distinct sets of metabolic genes were overrepresented in *sin-3(syb2172*) and *hda-3* datasets, and cell cycle genes in the *sin-3(syb2172*) dataset (Figure 3C). Interestingly SIN3 is associated with metabolic functions and cell cycle regulation in different species (Chaubal and Pile, 2018), suggesting possibly conserved regulatory functions. Neither components of GLP-1/Notch signaling, nor FBF proteins, key regulators of progenitor germ proliferation (Hubbard and Schedl, 2019) were found in our set of misregulated genes, while ribosomal protein genes were commonly downregulated in both mutants. *hda-3* upregulated gene were enriched in major sperm proteins and tau-tubulin kinases, which are both enriched in spermatogenic gonads . Comparison of our lists to those obtained by transcriptomic analyses of adult oogenic or spermatogenic gonads (Ortiz et al., 2014) confirmed that ‘spermatogenesis’ genes are overexpressed in *hda-3* mutants, and revealed that ‘oogenesis’ genes are overexpressed in *sin-3(syb2172*) mutants (Figure 3D).

To identify potential direct targets of SIN3 regulation, we compared our list of misregulated genes in *sin-3(syb2172*) mutants to a published list of SIN-3 binding sites obtained by ChIP-seq (Beurton et al., 2019). When considering all misexpressed genes, SIN-3 binding is observed on the promoter of both up- and down-regulated genes, with a small bias towards downregulated genes (Figure 3E). Interestingly however, among the set of SIN-3 target genes upregulated in *sin-3(syb2172*) mutants (146), near 50% of these are oogenic (Figure 3F). Altogether, our data suggest that HDA-3 represses spermatogenic genes, while SIN-3 may play a direct role in repressing oogenic genes, independently of HDA-3.

### SIN-3 contributes to silencing of the silenced X chromosome in the germline

Further analysis revealed that a large number (109) of genes upregulated in *sin-3(syb2172*) mutants reside on the X chromosome (Figure 4A), which represents a significant enrichment and suggests a role for SIN-3 in the repression of X-linked transcripts not shared with HDA-3 (Kelly et al., 2002). In *C. elegans*, the X chromosomes in XX hermaphrodites and XO males are globally “silenced” during most stages of *C. elegans* germ cell development through the combined activity of the MES (maternal effect sterile) proteins. MES-2/3/6 are homologs of PRC2 (Gaydos et al., 2014; Bender et al., 2006; Snel et al., 2022) and deposit repressive H3K27me3 (Bender et al., 2006; Gaydos et al., 2014). MES-4 instead deposits the activating mark H3K36me3 on germline expressed autosomal genes and antagonizes H3K27me3, leading to its concentration on the X (Gaydos et al., 2014; Bender et al., 2006). MRG-1, a component of several chromatin complexes in addition to SIN3S (Chen et al., 2010; Huang et al., 2017; Smith et al., 2013; Yochum and Ayer, 2002; Baytek et al., 2022), also contributes to silencing of the X in the *C. elegans* germline (Takasaki et al., 2007). Comparison of our list of *sin-3(syb2172*) upregulated genes on the X with a list of MES-3 and MRG-1 targets on the X (Cockrum and Strome, 2022) revealed commonly upregulated genes in the three mutants (37), as well as a larger set genes uniquely in common between *mes-3* and *mrg-1,* as previously described (Figure 4B) (Cockrum and Strome, 2022). In addition, 48 genes were upregulated in *sin-3* mutants only: these may be unique targets of SIN3 on the X (Figure 4B). We also observed enrichment of X linked genes among the small set of genes commonly upregulated in *hda-3* and *sin-3* mutants (Figure S5A), suggesting that SIN-3 may also repress a small set of genes in the context of SIN3L. One of these commonly upregulated genes is *lin-15B*, encoding a THAP domain transcription factor also upregulated in *mes-4* mutants. Loss of *lin-15B* in *mes-4* mutants was shown to reduces X misexpression and prevent germline death (Cockrum and Strome, 2022). We found that *lin-15B*(RNAi) had no effect on the fertility of *sin-3* mutants (Figure S5B), while its upregulation in *hda-3* mutants is not associated with any obvious germline defects, suggesting that additional mechanisms contribute to sterility in the absence of *sin-3*.

**Figure 4.**
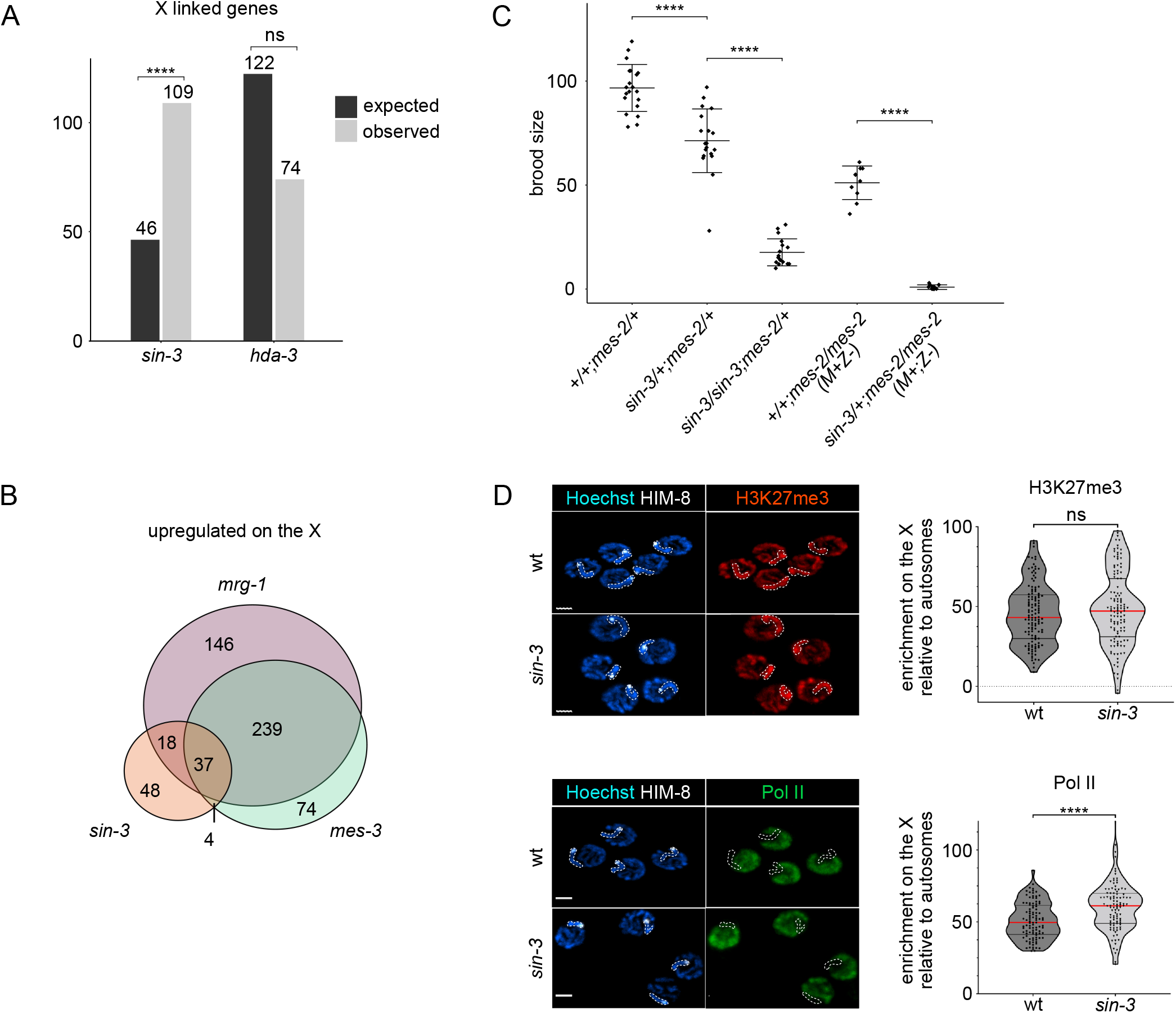
SIN-3 contributes to X chromosome silencing in the germline. (A) Enrichment analysis for upregulated genes in *sin-3(syb2172)* and *hda-3(ok1991)* mutants. Bar plot represent the expected (black) and observed (gray) number of upregulated X-linked genes in each mutant background. Enrichment was calculated with hypergeometric tests performed in R. *** p-value < 0,0001. Gene numbers in each set are the following: total number of analyzed genes for *sin-3* (7757), total number of genes on X chromosome (886), total number of *sin-3* up genes (405), number of *sin-3* up genes on X chromosome (109), total number of analyzed genes for *hda-3* (8407), total number of genes on X chromosome (831), total number of *hda-3* up genes (1238), number of *hda-3* up genes on X chromosome (74). (B) Venn Diagram showing overlap between X-linked genes upregulated in *sin-3(syb2172), mrg-1(qa6200)* and *mes-3(bn199)* mutants. *mrg-1* and *mes-3* gene lists are from (Cockrum and Strome, 2022). (C) Decreased dosage of wildtype *sin-3* enhances *mes-2* sterility. The total number of F1 progeny from animals with the indicated genotype was scored. M+Z-refers to homozygote *mes-2* mutants (Z-) with maternal contribution (M+). Multiple comparison was done using a generalized linear model with a quasi-Poisson law. **** p < 0.0001. (D) Representative confocal microscopy images of H3K27me3, pol II and HIM-8 staining on pachytene stage nuclei in wildtype and *sin-3(syb2172)* animals. HIM-8 staining identifies the X chromosome (outlined by dotted line). Scale bar is 2 µm. Violin plots display the enrichment on the X relative to autosomes for either H3K27me3 or pol II staining.

### Loss of *sin-3* enhances sterility of *mes-2*/PRC2

Altogether our data shows that the absence of *sin-3* results in maternal effect sterility, reduction in progenitor cell proliferation, and derepression of X linked genes. Although less severe, these phenotypes are reminiscent of those found in *mes* mutants (Capowski et al., 1991), and suggest that *sin-3* and *mes* genes may genetically interact. *mes* mutants are fully penetrant maternal effect sterile: homozygous (*mes/mes*) mutant offspring from heterozygous (*mes/+*) parents are fertile due to maternal rescue, but their offspring (M-Z-) are sterile. We compared the fertility of *+/+; mes-2/mes-2* and *sin-3/+*; *mes-2/mes-2* animals derived from *sin-3*(*syb2172*)*/+; mes-2(bn11)/+* mothers. As expected, *+/+ ; mes-2/mes-2* animals that have inherited maternal MES-2 protein were fertile (Figure 4C) (Capowski et al., 1991). *sin-3/+* ; *mes-2/mes-2* animals were instead fully sterile, showing that decreased *sin-3* dosage in these animals enhances the *mes-2* phenotype, despite their inheritance of maternal MES-2 protein. The effect of decreased *sin-3* dosage on fertility was also observed in *sin-3/+; mes-2/+* animals, whose brood size was reduced compared to their *+/+ ; mes-2/+* siblings, and further reduced in *sin-3/sin-3 ; mes-2/+* animals. Together, these results are consistent with SIN-3 and MES-2 playing similar functions in the germline, most likely through the silencing of distinct X-linked genes (Figure 4C), and reveal *sin-3* sensitivity to dosage.

MES proteins silence the X by concentrating repressive H3K27me3 on this chromosome (Bender et al., 2006). To test whether derepression of the X in *sin-3* mutants and enhancement of the *mes-2* phenotype following reduced dosage of *sin-3* is accompanied by a decrease in H3K27me3 on the X, we carried out immunostaining experiments with H3K27me3 specific antibodies on pachytene nuclei, when the X is normally silenced (Figure 4D). Using HIM-8 staining to identify the silenced X (Phillips et al., 2005; Rappaport et al., 2021), we observed that in wild type H3K27me3 was enriched on this chromosome with respect to autosomes, as expected (Kelly et al., 2002), and that this enrichment was still observed in *sin-3(syb2172*) mutants. Polymerase II (pol II) staining instead revealed a slight but reproducible increase on the X in *sin-3* mutant germlines, consistent with increased genes expression from this chromosome. Interestingly, in both H3K27me3 and pol II immunostaining experiments we observed a small subpopulation of nuclei with decreased H3K27me3 or increased pol II signal in *sin-3* mutants compared to wiltype (Figure 4D). Our experimental setup did not allow us to establish whether these are the same nuclei, that is, whether nuclei with decreased H3K27me3 also show increased pol II staining. Nonetheless, these results suggest that within a single *sin-3* mutant germline, the chromatin and transcriptional state of the X may vary between individual pachytene nuclei.

### SIN-3 inactivation results in precocious and stochastic transcription of autosomal and X-linked genes

To gain further insight on how loss of *sin-3* affects transcription at the single cell level, we performed smiFISH experiments (Tsanov et al., 2016) with exonic probes for the X-linked genes *lin-15B* and *nmy-1,* both of which are upregulated in *sin-3(syb2172*) mutant germlines (Table S2). In wildtype germlines, both transcripts were mostly detected as spots starting in late pachytene, consistent with the re-expression of X-linked genes at this stage (Kelly et al., 2002; Tzur et al., 2018) (Figure 5). The occasional spots observed more distally likely represent the few transcriptional events that take place at earlier stages. In *sin-3* mutant germlines we observed a significant increase in the number of detected transcripts starting in the transition zone/early pachytene, suggesting precocious re-expression of X-linked genes. In addition, the accumulation of fluorescent spots was highly heterogeneous, with groups of cells containing many spots found immediately adjacent to nuclei with only a few or no spots (Figure 5). Therefore, in *sin-3* mutants, re-expression of X linked transcripts occurs in a stochastic manner within a single germline.

**Figure 5.**
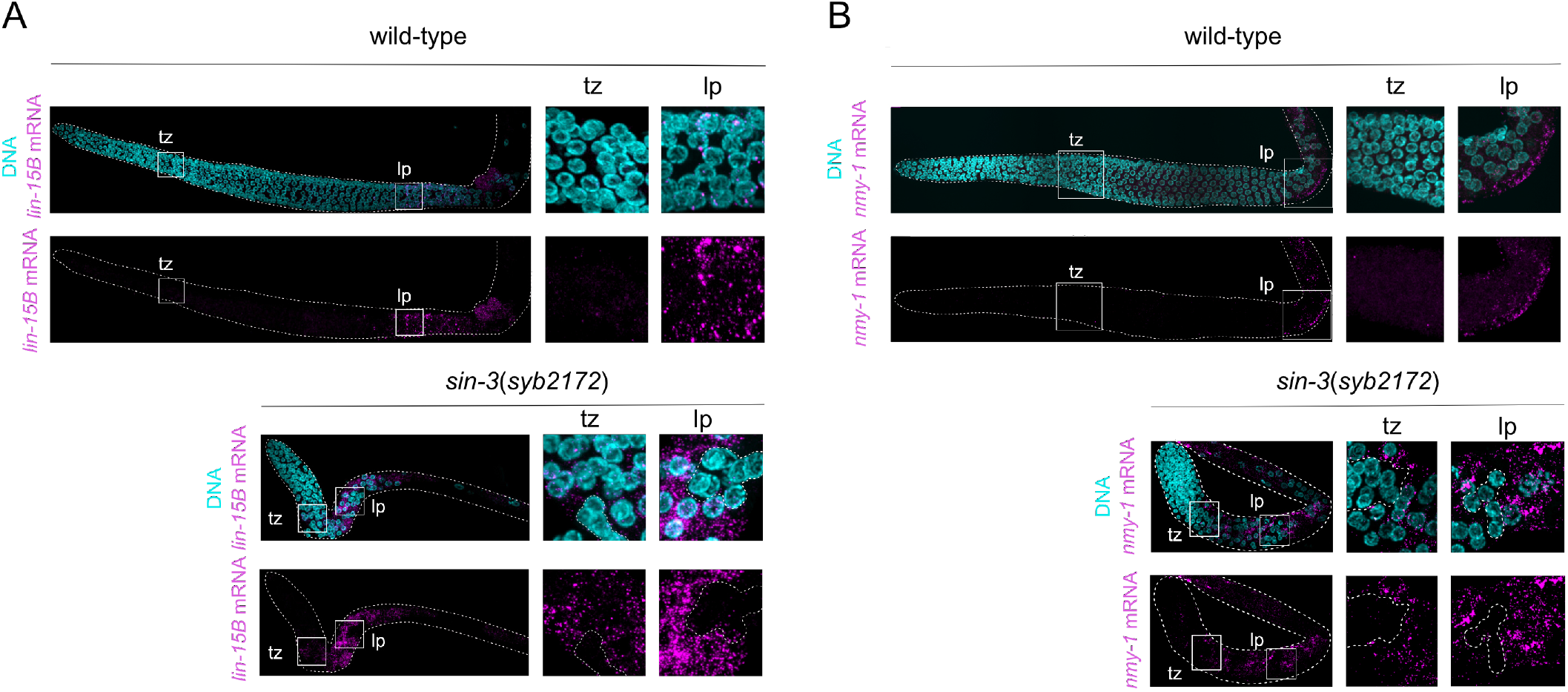
*sin-3* germ nuclei precociously and stochastically desilence X-linked genes. Images show expression of *lin-15B* (A) and *nmy-1* (B) X-linked transcripts detected by smFISH analysis in wildtype and *sin-3*(*syb2172*) germlines. Transcripts are detected as bright nuclear or cytoplasmic dots (magenta). A zoom of the boxed regions representing nuclei from transition zone (tz) and late pachytene (lp) are presented on the right. Transition zone nuclei are defined by crescent shaped morphology, and pachytene nuclei by the typical “bowl of spaghetti” appearance observed by Hoechst staining (cyan) (Crittenden et al., 2017). White broken lines outline examples of group of nuclei with few or no transcripts neighboring nuclei with strong expression. Images are maximum Z projections.

### Increased acetylation on specific residues in *sin-3* mutants

SIN-3 alone does not have catalytic activity, but is closely associated with class I HDACs that target histones or proteins for deacetylation (Kelly and Cowley, 2013; Laherty et al., 1997; Milazzo et al., 2020). To identify histone modifications potentially targeted by SIN-3 and its associated HDACs, we carried out MS analysis of purified histones from *sin-3(tm1276)* young adults, a stage at which the germline is fully developed. We chose the *tm1276* allele because the sterility of *syb2172* does not allow the culturing of a large enough number of animals required for this type of analysis. While the abundance of most quantified histone acetylations was not significantly altered in *sin-3(tm1276)* mutants, we detected a significant increase in the bivalent mark K18AcK23Ac on histone H3, and in K27Ac and K27me1 on histone H3.3 (Figure 6A). The abundance of histone H3 with acetylation on either K18 or K23 residues alone was not significantly affected. Coexistence and positive interplay between K18Ac and K23Ac has previously been described (Klein et al., 2019; Schwämmle et al., 2014). Semi-quantitative western-blot analysis using antibodies that detect acetylation marks on both histone H3 and H3.3 confirmed increased acetylation on K27 and K18, but not K9 in total extracts from *sin-3(tm1276)* mutant young adults (Figure 6B). No change in acetylation levels was observed in *hda-3* mutant extracts, suggesting that another SIN-3 associated HDAC, most likely HDA-1, is responsible for deacetylation of these residues. ChIP-qPCR analysis on a set of genes misregulated in both *sin-3* mutant alleles showed no clear correlation between acetylation levels and mRNA levels: we observed both increased and decreased H3K27Ac on the promoter of genes upregulated in *sin-3* mutants (Figure S7A). This lack of correlation may reflect heterogeneity in the population of germ cells, as indicated by the smiFISH analysis. Increased H3K27Ac in the germline was further confirmed in immunostaining experiments on *sin-3(tm1276)* animals using H3K27Ac antibodies (Figure S7B).

**Figure 6.**
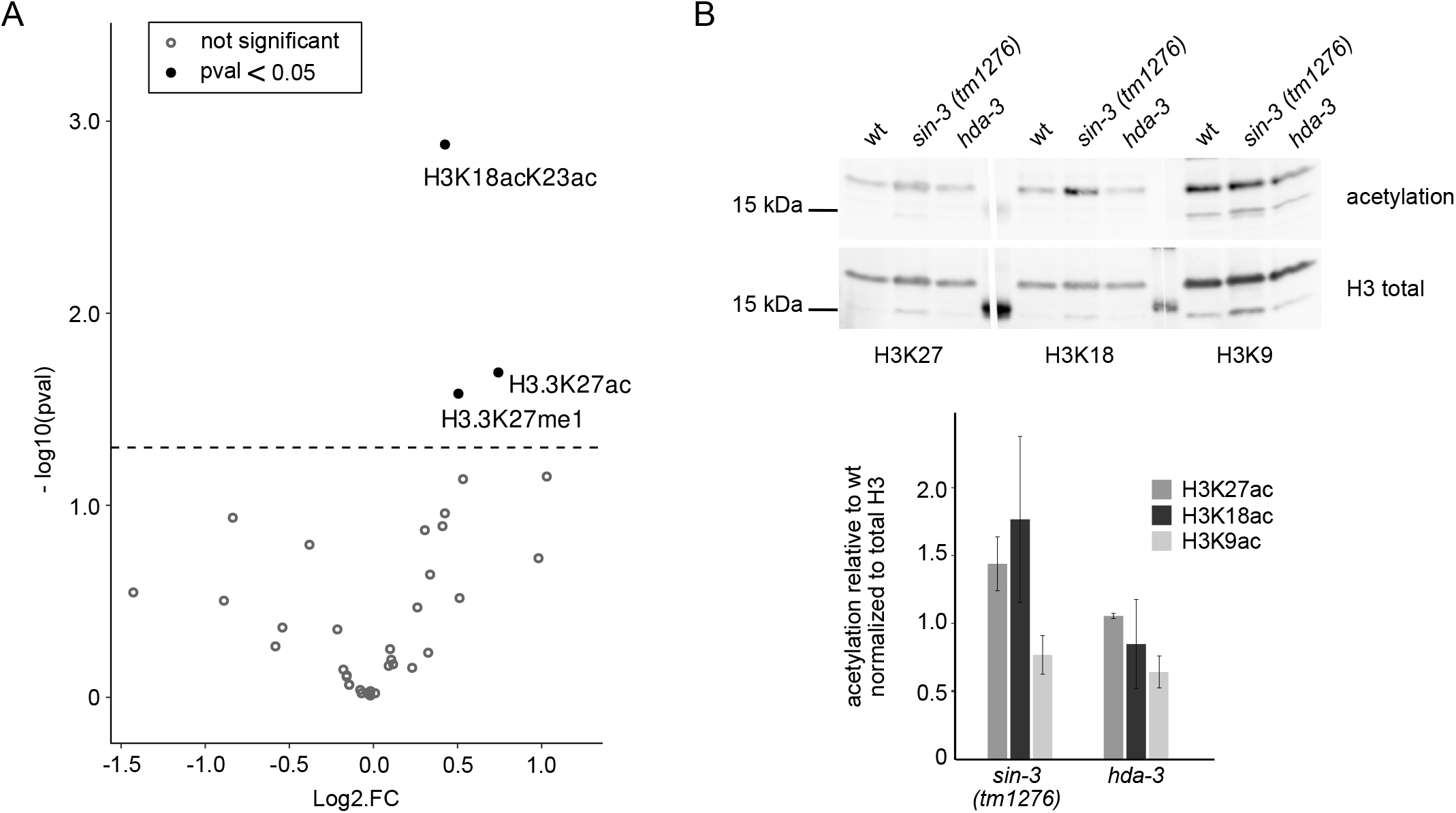
Specific residues in *sin-3(tm1276)* mutants are more acetylated than in wt. (A) Volcano plot representing the log2 fold change (log2FC) on H2, H3 and H4 peptide modification abundances in *sin-3(tm1276)* versus wildtype young adult worms plotted against statistical significance (log10(p-value)). Modifications that are significantly different between both genotypes are plotted as red dots. Horizontal blue dashed line represents a significant threshold of p-value < 0.05. (B) Semi-quantitative western blot analysis of acetylation levels in total extracts from wt, *sin-3(tm1276)* and *hda-3(ok1991)* young adults. Acetylation of H3K9, H3K18 and H3K27 was detected by fluorescence after western blot using specific antibodies. Levels of each modification were quantified with Image J and normalized to total H3 levels. Barplot represented the mean of two independent biological replicates; error bars correspond to SD.

## Discussion

In this study we have shown that the highly conserved SIN3 transcriptional coregulator is essential for fertility in *C. elegans*. We biochemically identified SIN-3 interactors that differentially contribute to fertility and provide data suggesting that specific class I HDACs may reside in distinct complexes. We further uncovered a role for SIN-3 in the timely repression of X-linked genes, and showed by single cell analysis that re-expression of X-linked genes in *sin-3* mutant germlines occurs prematurely and in a stochastic manner. These results suggest that the absence of SIN-3 may not uniformly affect gene expression in all germ cells.

Maternal effects sterility associated with *sin-3* inactivation was revealed here using a CRISPR-Cas9 loss-of-function allele*, syb2172,* and was missed in previous studies using the hypomorphic allele *sin-3(tm1273*). Sterility in *sin-3*(*syb2172)* is associated with reduced proliferation of progenitor germ cells, a process tightly controlled by GLP-1/Notch signaling and RNA regulators (Hubbard and Schedl, 2019). However, expression of these factors is not affected in *sin-3* mutant germlines, nor did we find any evidence for defects in the switch between self-renewal and differentiation that could account for a reduced pool of progenitor cells <u>(Crittenden et al. 2017)</u>. What we instead observed is increased expression of X-linked genes. Silencing of the X in the germline is under the control of MES proteins; in *mes* mutants upregulation of X linked genes results in death of nascent germ cells, a more severe phenotype than observed in the absence of *sin-3* (Bender et al., 2006; Gaydos et al., 2014). This may be partly accounted for by the significantly smaller number of SIN-3 target genes on the X. A critical target for repression by MES proteins on the X is the *lin-15B* THAP domain transcription factor, and its inactivation prevents germline death in *mes* mutants (Cockrum and Strome, 2022). While *lin-15B* is also upregulated in both *sin-3* and *hda-3* mutant germlines, its inactivation in *sin-3* mutants had no effect on fertility, suggesting that in this context its upregulation alone does not contribute to sterility. Consistent with additional SIN-3 functions in the germline, transcriptomics analysis showed that SIN-3 regulates genes with roles in metabolism and cell cycle regulation. Additional roles for SIN-3 independent of MES, either on the X itself or at other loci, are supported by genetic analysis in which we observed that loss of a single copy of *sin-3* enhanced the sterility of *mes-2* M+Z-mutants with maternal MES-2 contribution.

Our analysis also provided important insight into the composition and function of distinct SIN3 associated complexes in a single tissue. We confirm previous results showing that knock-down of SIN3S complex subunit ATHP-1/PHF12 results in reduced fertility (Beurton et al., 2019), and show that animals lacking SUDS-3, an essential component of SIN3L complexes (Banks et al., 2020; Clark et al., 2015), are instead fully fertile. Like SIN-3, SIN3S component MRG-1 is required for silencing of the X chromosome and fertility (Takasaki et al., 2007), and both SIN-3 and MRG-1 are required for piRNA silencing in the germline (Kim et al., 2021). These shared phenotypes are consistent with common functions as part of the same complex, although MRG-1 most likely has additional functions alone or in the context of additional complexes (Baytek et al., 2022; Hajduskova et al., 2018; Bleuyard et al., 2017; Iwamori et al., 2016; Iwamori et al., 2016; Smith et al., 2013). Although SIN3L subunits alone may not be required for fertility, the observation that *athp-1; hda-3* and *athp-1; suds-3* double mutants are fully sterile suggests at least partial redundancy of the two complexes, or additional functions for the encoded subunits independent of SIN-3.

Interestingly, while mammalian HDAC1 and HDAC2 are mostly considered to be functionally redundant, we found that both HDA-3 and HDA-1 are present in the SIN3L complex identified in this study, while SIN3S components only copurify with HDA-1(Baytek et al., 2022; Beurton et al., 2019). While HDA-1 has been found in at least two additional complexes, NuRD and MEC (Passannante et al., 2010; Kim et al., 2021), an HDA-3 containing complex has not been described to date. Although mammalian HDAC1 and 2 copurify in mammalian SIN3 complexes (Banks et al., 2018; Banks et al., 2020), they also have unique functions and partners (Terzi Cizmecioglu et al., 2020; Quaas et al., 2022). Notably, in mESCs ARID4B was found to interact with SIN3A and HDAC1, but not HDAC2 (Terzi Cizmecioglu et al., 2020). Interestingly the mammalian counterpart of transcription factor TCF-19, identified here as an accessory component of SIN3L, was found to interact with NuRD/HDAC at the promoter of gluconeogenic genes (Sen et al., 2017), suggesting that it may play a role in the recruitment of distinct HDAC containing complexes to chromatin. Consistent with unique functions, HDAC1 knockout mice are embryonic lethal (Lagger, 2002), whereas an HDAC2 deletion is viable (Trivedi et al., 2007). Similarly *hda-1* inactivation results in larval lethality, while *hda-3* mutant are viable (Kawamura and Maruyama, 2020 and this work).

Our expression profiling shows that, as in other systems, loss of SIN-3 results in both repression and activation of gene expression (Saha et al., 2016; Saunders et al., 2017; Yao et al., 2017; van Oevelen et al., 2010), consistent with the presence of HDACs at promoters of actively transcribed as well as repressed genes (Wang et al., 2009). SIN-3 may influence gene expression in several ways. On the silent X chromosome, we did not observe depletion of H3K27me3 in the absence of SIN-3, arguing against a simple model in which desilencing results from loss of repressive chromatin. Immunostaining of *sin-3* mutants showed a small increase in pol II on the X in pachytene, while smFISH analysis revealed that X-linked transcripts accumulate stochastically in early meiotic cells, a stage in which the X chromosome is normally silenced (Kelly et al., 2002). Together, these results suggest that SIN-3 may not elicit a uniform response on its targets on the X, but instead act as an “epigenetic switch” at individual target genes, resulting in ON or OFF transcriptional states. This may occur in a concerted manner, or independently at each locus. Perturbation of lysine acetylation is one mechanism that could potentially modify the frequency of transcriptional bursting, or switching between states (Nicolas et al., 2018; Tunnacliffe and Chubb, 2020), but whether SIN3 influences gene expression as specific loci through HDAC activity remains to be established. We identified H3K18AcK23Ac as a SIN-3 dependent bivalent mark, and confirmed increased H3K18Ac in total extracts from *sin-3(tm1276)* mutant animals. Interestingly, although a partially reconstituted SIN3B complex displayed very low deacetylase activity at every H3 acetyl-Lys site tested in *in vitro* assays on reconstituted nucleosomes, a slight preference for H3K18Ac, H3K23Ac and H3K27Ac compared to K9Ac was observed (Wang et al., 2020). A preference for binding to diacetyladed residues has been reported for several readers of acetylation (Parthun, 2012; Barman et al., 2021; Obi et al., 2020), with one of the two bromodomain of BRDT showing highest affinity for H3K18AcK23Ac (Miller et al., 2016).

Although ChIP-qPCR experiments of a number of hand-picked genes did not reveal a correlation between increased H3K27Ac and expression levels in *sin-3* mutants, stochastic effects of SIN-3 depletion on gene expression, as we observed on the X, may require single cell approaches to further address this question. In addition, SIN-3 may play non-canonical roles in repressive chromatin (Torres-Campana et al., 2022), target non-histone substrates (Milazzo et al., 2020), or have functions independent of HDAC catalytic activity, including nucleosome assembly (Chen et al., 2012), as suggested by the physical interaction with the NAP1 chaperone (Moshkin et al., 2009 and our data). Some of these activities may also contribute to SIN-3 mediated gene activation.

The dosage effect of *sin-3* inactivation on the *mes* phenotype is particularly interesting given that SIN-3 haploinsufficiency is the underlying cause of Witteveen-Kolk syndrome and related intellectual disability (ID)/autism syndromes (Balasubramanian et al., 2021; Latypova et al., 2021; Witteveen et al., 2016). Future studies in simpler systems such as the *C. elegans* germline may help gain insight into common regulatory mechanisms involving SIN3 and aid in the identification and study of pathogenic SIN3 variants associated with disease.

## Materials and Methods

### Strains and maintenance

Nematode strain maintenance was as described previously (Brenner, 1974). wild type N2 (Bristol) was used as reference. Strains used are as follows (* from (Beurton et al., 2019); **created for this study): *PFR590, *sin-3(tm1276)* I; **PHX2172*, sin-3(syb2172)/hT2[bli-4(e937) let-?(q782) qIs48] I*; *PFR588, *cfp-1(tm6369)* IV; *PFR593, *athp-1(tm4223) III*, **PHX2212, *suds-3(syb2212)* V; *PFR740, *hda-3(ok1991) I*; **PFR746, *athp-1(tm4223)*III; *hda-3(ok1991)*I; **PFR747, *athp-1(tm4223)* III; *suds-3(syb2212)*V; **PHX4769, *arid-1(syb4769)* V; **PHX5132, *hda-3(syb5132) I*; SS186, *mes-2(bn11)*; *unc-4(e120)/mnC1[dpy-10(e128) unc-52(e444)]* II. CRISPR-Cas9 alleles were created by SunyBiotech using the primers listed in Table S3.

### Genetic Crosses

Strains carrying specific combination of mutations, transgenes and CRISPR-Cas9 knock-in alleles were obtained by crossing. Males used for crosses were either heterozygotes obtained from crossing wildtype males with hermaphrodites carrying the mutation or transgene of interest, or homozygotes obtained by heat shock of mutant L4 hermaphrodites. Single hermaphrodites from crosses were isolated and allowed to lay eggs overnight, then genotyped by PCR and/or screened for the desired phenotype. This was repeated until double or triple homozygotes were obtained.

### Scoring brood size

10 L4 worms per genotype were isolated onto single NGM plates seeded with OP50 bacteria, allowed to grow into egg-laying adults overnight at 20°C, then transferred every 12 hours to fresh plates until they ceased laying eggs. Plates were then scored for the number of viable progeny and dead embryos that failed to hatch 24 hours after removal of the mother.

### Observation of live worms

Worms were placed on 2% agarose pads in a 15µL drop of 10mM Levamisole in M9. A cover slip was placed on the sample, and slides observed using either brightfield, fluorescence or differential interference contrast (DIC) microscopy with a Zeiss Axio Imager A2 microscope or Nikon AZ100M Zoom microscope.

### Immunofluorescence microscopy on dissected gonads

Experiments were as previously described (Herbette et al., 2017). Briefly, young adults (L4 + 12h) were transferred to an empty plate for 15 min to remove bacteria, then transferred to a drop of dissection buffer (M9 0,4X, levamisole 10mM) on polylysine coated slides slides. Worms were cut at the pharynx and gonads extracted using 30G x ½” needles (BD Microlance). Dissected gonads were fixed in 3,2 % PFA (ThermoFisher) for 5 min, and placed on dry ice prior to freeze-cracking (Strome and Wood, 1982). Slides were immersed in MeOH at −20 °C for 1 min, washed 3 times in PBS 1X (Sigma) with 0,1% Tween20 (Sigma), blocked for 40 min in PBS 1X, 0.1% Tween20 (PBST) and 1% BSA (MP Bio), and incubated with primary antibodies overnight at 4 °C in a humid chamber. Slides were washed 3 times in PBST, incubated with secondary antibodies for 50 min, then washed 7 min in PBST plus Hoechst (Sigma) 5 μg/ml, 2 times 10 min in PBST, and mounted in mounting medium (PBS 1X, 90% glycerol, 0.4% propyl gallate). All antibodies were diluted in 1X PBS, 0.1% Tween 20, 1% BSA. Antibodies used: mouse anti-H3K37me3 (#61017 Active Motif, 1:300), rabbit anti-H3K27ac (#39133 Active Motif, 1:5000), mouse anti-H3 (#14269 Cell Signaling, 1:500), Rat anti-RNA Pol II (#04-1571 Millipore, 1:1000), rabbit anti-pH3 (#SC-8656-R Santa Cruz Biotechnology 1:100), rabbit anti-HIM-3 (Kind gift from Monique Zetka), rat anti-HIM-8 (Kind gift from Abby Dernburg (UC Berkeley), goat anti-rat (#21434 Invitrogen, 1:1000), goat anti-mouse Alexa Fluor Plus 555 (#A32727 Invitrogen 1:1,000), goat anti-rabbit Alexa Fluor Plus 647 (#A32733 Invitrogen 1:1,000). Antibody dilutions were in PBST. Images were acquired with either a Zeiss LSM980 confocal microscope or a Yokogawa CQ1 spinning disk confocal microscope.

### Measuring signal ratio between X and autosomes

Immunofluorescent acquisitions of late pachytene nuclei obtained with a Zeiss LSM980 confocal microscope were analyzed using FIJI. Maximum projections of 4 to 6 stacks of 0.16 µm were obtained, and nuclei where both X chromosome (identified using HIM-8 staining) and at least one autosome were present in the same transversal position were analyzed in subsequent steps. The average intensity on the X and one autosome was quantified, background intensity subtracted and the ratio between the X and autosome was then quantified on each nucleus analyzed. 100 nuclei were analyzed per genotype. Statistical analysis was performed using t-tests.

### Acridine Orange (AO) staining of apoptotic cells

Staining of apoptotic cells was as previously described (Gumienny et al., 1999). 1 mL of 50µg/mL Acridine Orange (AO, Sigma) in M9 1X was dropped on plates containing 20 to 30 young adults (L4 + 12h) for each genotype. Plates were kept at 20°C for 2 hours in the dark, and worms then transferred onto fresh plates to wash excess AO. After washing, worms were individually transferred onto agarose pads and imaged using Zeiss LSM980 Confocal Microscope. Images were processed using FIJI (Schindelin et al., 2012), and cells positives for AO staining were counted in z-stack. 20 gonads per genotypes were counted.

### Immunoprecipitation experiments

Immunoprecipitations were performed on frozen embryos prepared by hypochlorite treatment from animals grown at 20°C on enriched NGM seeded with 10X concentrated HB101 bacteria, as previously described (Beurton et al., 2019). After hypochlorite treatment, embryos were washed once in IP buffer (50 mM HEPES/KOH, pH 7.5; 300 mM KCl; 1 mM EDTA; 1 mM MgCl_2_; 0.2% Igepal-CA630; and 10% glycerol) and flash-frozen in beads in liquid nitrogen. Embryos were then grounded to powder, resuspended in 1 bead volume of IP buffer containing 2X complete protease inhibitors (Roche) and sonicated on ice at an amplitude of 30% for 2.5 min (15’’ ON/15’’ OFF pulses) using an Ultrasonic Processor (Bioblock Scientific). Protein extracts were recovered in the supernatant after centrifugation at 20,000*g* for 15 min at 4°C. Protein concentration was estimated using the Bradford assay (Bio-Rad Protein Assay Dye). All immunoprecipitations were performed with 70 mg of total protein extract in 10 ml diluted in IP buffer. Each sample was incubated for preclearing with 200 μL slurry of binding control magnetic agarose beads (ChromoTek bmab) for 1 h at 4°C. Then 200 μL of GFP-TRAP MA (ChromoTek gtma) or 300 μL RFP-TRAP MA beads slurry (ChromoTek rtma) were added to the sample. Beads incubation was performed 3h on a rotator at 4°C. Beads were collected with a magnet, washed three times in IP buffer and once in Benzo buffer (HEPES/KOH 50 mM, pH 7.5; KCl 150 mM; EDTA 1 mM; MgCl_2_ 1 mM; Igepal-CA630 0.2%; and glycerol 10%). Beads were then incubated in 400 μl of Benzo buffer containing 2,500 units of benzonase (Sigma-Aldrich) for 1 h at 4°C and washed three times in IP buffer. Eluates were recovered by incubation at 95°C for 10 min in 60 μl of 1× LDS buffer (Thermo Fisher Scientific). 1/10 of each eluate was resolved on a 4–12% NuPage Novex gel (Thermo Fisher Scientific) and stained with SilverQuest staining kit (Thermo Fisher Scientific). 40 μl of the eluates was then analyzed by mass spectrometry.

### MS-based proteomic analyses of IP eluates

Proteins from IP eluates solubilized in Laemmli buffer were stacked in the top of a 4-12% NuPAGE gel (Invitrogen), stained with Coomassie blue R-250 (Bio-Rad) before in-gel digestion using modified trypsin (Promega, sequencing grade) as previously described (Casabona et al., 2013). The resulting peptides were analyzed by online nanoliquid chromatography coupled to MS/MS (Ultimate 3000 RSLCnano and Q-Exactive HF, Thermo Fisher Scientific) using 120 min and 90 min acetonitrile gradients for SIN-3 and ARID-1 interactomes respectively. For this purpose, the peptides were sampled on a precolumn (300 μm x 5 mm PepMap C18, Thermo Scientific) and separated in a 75 μm x 250 mm C18 column (Reprosil-Pur 120 C18-AQ, 1.9 μm, Dr. Maisch). The MS and MS/MS data were acquired using Xcalibur 4.0 (Thermo Fisher Scientific).

Peptides and proteins were identified by Mascot (version 2.8.0, Matrix Science) through concomitant searches against the Uniprot database (*Caenorhabditis elegans* taxonomy, 20220531 download), a homemade database containing the sequences of the bait proteins, and a homemade database containing the sequences of classical contaminant proteins found in proteomic analyses (human keratins, trypsin…). Trypsin/P was chosen as the enzyme and two missed cleavages were allowed. Precursor and fragment mass error tolerances were set at respectively at 10 and 20 ppm. Peptide modifications allowed during the search were: Carbamidomethyl (C, fixed), Acetyl (Protein N-term, variable) and Oxidation (M, variable). The Proline software (Bouyssié et al., 2020), version 2.2.0) was used for the compilation, grouping, and filtering of the results: conservation of rank 1 peptides, peptide length ≥ 6 amino acids, false discovery rate of peptide-spectrum-match identifications < 1% (Couté et al., 2020), and minimum of one specific peptide per identified protein group. MS data have been deposited to the ProteomeXchange Consortium via the PRIDE partner repository (Perez-Riverol et al., 2019) with the dataset identifier PXD0XXXXX. Proline was then used to perform a spectral counts-based comparison of the protein groups identified in the different samples. Proteins from the contaminant database were discarded from the final list of identified proteins. To be considered as a potential binding partner of a bait, a protein must be detected with a minimum of three specific spectral counts, identified only in the positive eluate, or enriched at least three times in this eluate compared to the corresponding control eluate on the basis of spectral counts.

### RNA-sequencing by Smart Seq and data analysis

For each biological replicate, gonads from 9-12 young adults were dissected in UltraPure water on microscope slides as described for immunofluorescence, except that the entire dissection protocol was performed at 4°C to immobilize worms without use of anesthetic RNA was isolated, reverse-transcribed, amplified and clean up following the Smart-seq2 protocol described by Serra et al. (Serra et al., 2018). Three independent biological replicates were performed for each strain. Librairies were generated at the GenomEast Platform [IGBMC, Strasbourg, France] using the SMART-Seq v4 UltraLow Input RNA kit (Clontech) followed by the Nextera XT DNA sample preparation Kit (Illumina) and sequenced using the Illumina Hiseq 4000 technology (1x50 bases). For data analysis, fastq files were processed with fastp (Version 0.20.1), reads were mapped to the *C. elegans* reference genome (WS278) by RNA-STAR (Version 2.7.3a). and gene expression level in each sample was calculated by htseq-count (Version 0.7.2). Differential expression between each mutant strain and wild type was then calculated with DESeq2 (Version 1.36.0) using a homemade R script (Rversion 4.2.2). At this step PCA reveals that one replicate for *hda-3* mutant was an outlier and thus it was discarded for further analysis. Of the 46934 annotated genes, only genes with a baseMean > 10 were selected in each condition (8407 genes for *hda-3* and 7757 for *sin-3*). Gene expression data are available at GEO with the accession XXXXX

### mRNA detection by smiFISH

smFISH was performed using a smiFISH protcol provided by the Hubstenberger lab and adapted from 10.1093/nar/gkw784. Primary probes were designed using the R script Oligostan (https://bitbucket.org/muellerflorian/fish_quant/src/master/Oligostan/; (Tsanov et al., 2016). *lin-15B* and *nmy-1* primary probes were designed with rules 1, 2, 3, 4, and 5 in the PNASfilterOption and *C04F12.1* and *cdc-14* primary probes were designed with rules 1, 2, 4 and 5. For all probes, MinGC content was set up at 0 .4 and maxGC content at 0.6. Primary probes sequences are provided in Table S4. Probes were produced by hybridizing of primary probes with a FLAPx-Cy5 secondary probe in TE 100 mM NaCl. Denaturation was performed at 95°C for 5 minutes and the mix was allowed to cool down at room temperature until it reaches 35-40°C. Worms were dissected for gonad isolation on Poly-Lysine coated slides in PB buffer complemented with Levamisole. Gonads were pre-fixed in 4% paraformalhéhyde for 4 minutes at room temperature and immediately frozen on dry ice. After freeze-cracking, gonads were fixed in cold PFA 4% for 20 minutes at room temperature and washed twice with PB buffer before a 20 hour fixation at 4°C in cold 70% EtOH. Before hybridization, slides were washed twice in 1X PBS and once in 1X SSC complemented with 15% of formamide. An addition 20 minutes incubation was perfomed for equilibration in 1X PBS/ 15% formamide before proceeding with hybridization. For hybridization, probes were diluted 40 times in Stellaris RNA hybridization buffer complemented with 10% formamide. Samples were incubated for at least 16 hours at 37°C. Samples were washed twice with 1x SSC complemented with 25% Formamide for 30 minutes and incubated with Hoechst for 5 minutes before two additional washes in 1X PBS. Samples were mounted in ProLong Antifade Mounting media (Invitrogen) and imaged using a Confocal AxioObserver Z1 LSM980 with AiryScan2 (Zeiss).

### Histone purification and Mass spectrometry analysis

Histones from wildtype and *sin-3(tm1276)* young adult worms were purified following Basic protocol 2 published by Millan-Ariño *et al* (Millan-Ariño et al., 2020). Mass spectrometry was then performed according to Basic Protocol 3 and histone peptides were analyzed as described in the same manuscript by bottom-up MS with a *C. elegans*−adapted version of the software EpiProfile 2.0 (Yuan et al., 2018). Raw data from Epiprofile were analyzed following recommendations from (Thomas et al., 2020) https://github.com/DenuLab/HistoneAnalysisWorkflow). Briefly, raw peptide abundance values were filtered to remove all modifications with missing values in more than one replicate for each genotype. Normalization of each modification was performed by dividing it by the sum of all modifications in the replicate. Finally, normalized values whose mean standard deviation is more than 60 % of its average were discarded. Statistical analysis was realized using the linear model : lm(normalized peptide abundance ∼ genotype) with a homemade script in R. All samples were analyzed in biological triplicate.

### Western blot analysis on histone marks

wildtype N2, *hda-3(ok1991)* and *sin-3(tm1276)* young adult worms were collected in M9 buffer, washed 3 times, pelleted and frozen in dry ice. After thawing, pellets were resuspended in TNET buffer (50 mM Tris·HCl (pH 8), 300 mM NaCl, 1 mM EDTA, 0,5% Triton X-100 and cOmplete™ Protease Inhibitor Cocktail (Merck # 11697498001])) and lysed with zirconium beads (Lysing Matrix Y, MP Biomedicals #116960050) using a Precellys 24 homogenizer (Ozyme)] with the following parameters : 6000 rpm 2 x 10 sec. Homogenates were centrifuged and supernatants aliquoted and frozen at −80◦ C. Total protein amount was quantified by the Bradford assay (Bio-Rad Protein Assay Dye).. 27 µg of protein extracts were loaded on 12% NuPage Novex gels for western blot analysis. After transfer, membranes were incubated overnight with the following antibodies diluted at 1:2,500: anti-H3 (clone 1B1B2) (Cell Signalling Technology, #14269]) anti-H3K9ac (Active Motif, #39137) anti-H3K27ac [Active Motif, #39133] and anti-H3K18ac antibody [Active Motif, #39755] and then for 1 hour with goat anti-rabbit DyLight™ 800 (Invitrogen # SA5-10036) and IRDye® 680RD goat anti-mouse (LiCOR #926-68070) diluted at 1:10,000. Aquisition was performed on a ChemiDoc MP apparatus (Bio-Rad). Quantification was performed carried out using Image J, and each acetylation signal was normalized to the level of histone H3. Two independent biological replicates were used for each strain for quantification.

## Acknowledgments

This work was supported by ANR (Agence Nationale de la Recherche) grant N° 19-CE12-0025-01 and the Centre National de la Recherche Scientifique. The proteomic experiments were partially supported by ANR grant ProFI (Proteomics French Infrastructure, ANR-10-INBS-08) and GRAL, a program from the Chemistry Biology Health (CBH) Graduate School of University Grenoble Alpes (ANR-17-EURE-0003). We thank Arnaud Hubstenberger for providing the smiFISH protocol before publication. We thank the Proteomics Biomedicum core facility at Karolinska Institutet for MS measurements. C.G.R. was supported by grants from the Swedish Research Council (VR) (2017-06088 and 2019-04868), the Swedish Cancer Society (Cancerfonden) (20 1034 Pj), and the Novo Nordisk Foundation (NNF21OC0070427 and NNF22OC0078353).

